# An experimentally verified mechanistic model for predicting quorum sensing-based switches

**DOI:** 10.1101/2025.06.05.658036

**Authors:** De Baets Jasmine, De Paepe Brecht, De Mey Marjan

**Author notes:** Corresponding author: M. De Mey, Coupure Links 653, 9000 Ghent, Belgium.

## Abstract

Quorum sensing-based genetic circuits are gaining traction in synthetic biology as they link population-level behaviour to individual cell responses. However, tuning these circuits remains challenging due to complex dynamics, particularly during the ‘Learn’ phase of the Design-Build-Test-Learn (DBTL) cycle. To accelerate this process, we developed a mathematical model to predict how varying expression levels of the transcription factor and synthase affect the response of the EsaI/EsaR quorum sensing system. A strain library was constructed, and experimental data were used to optimize the model. The final model could successfully differentiate between the effects of these expression levels on the response of the bidirectional promoter. It allowed visualization of all potential system outcomes and emphasized the transcription factor’s critical role in tuning the circuit. This model offers a valuable tool for fine-tuning EsaI/EsaR-based systems for synthetic biology applications. Moreover, given the homology within the LuxR-family quorum sensing systems, this modelling approach may serve as a foundation for model-based tuning of other quorum sensing systems.

## 1. Introduction

In nature, the production of costly public goods, such as virulence factors, bioluminescence and exopolysaccharides for biofilm formation, is controlled by a mechanism called quorum sensing (QS) (Schuster et al., 2013). The production of these goods places a high burden on the cells but is also very valuable for the population. Quorum sensing ensures that sufficient cells are present before production gets initiated so that the benefit of this public good for the population is higher than the burden caused by its production. The power of quorum sensing to connect processes occurring within the cell with the state of the population holds great potential for applications in the field of synthetic biology. It allows for the synchronization of response (Wu et al., 2020), synthetic communication between different bacteria in a consortium (Brenner et al., 2007; Dinh et al., 2020) and the regulation of biosynthetic pathways (Kim et al., 2020).

Acyl-homoserine lactone (AHL)-based quorum sensing systems, such as the LuxI/LuxR system, are especially easy to implement, since they only require the expression of the AHL-synthase and transcription factor to allow expression from the QS-regulated promoter. The synthase protein constitutively produces the AHLs, which can diffuse freely over the cell membrane. Therefore, at increasing cell densities, the AHL concentration will increase rapidly. Once a certain threshold concentration is reached, the transcription factor will be able to bind to it, leading to a change in promoter activity of the promoter it is regulating. Unlike most other transcription factors from the LuxR-family, EsaR binds a bidirectional promoter region (P_esaR/esaS_) in the absence of the AHL-molecule, 3-oxo-hexanoyl homoserine lactone (3OC6-HSL) (Schu et al., 2009). Furthermore, it functions as both a transcriptional activator and repressor of the P_esaS_ and P_esaR_ promoter, respectively (Figure 1). In the presence of 3OC6-HSL, at high cell densities, EsaR will undergo a conformational change and is, therefore, no longer capable of binding the bidirectional P_esaR/esaS_ promoter (Schu et al., 2009). Because of its dual functionality, this system allows for the creation of more complex circuitry or the simultaneous up- and downregulation of different pathways. The latter was applied for the production of poly-β-hydroxybutyrate and 5-aminovulinic acid, where the production pathway was upregulated and the competing pathways downregulated once a certain cell density was reached to reduce competition between growth and production (Gu et al., 2020).

**Figure 1.**
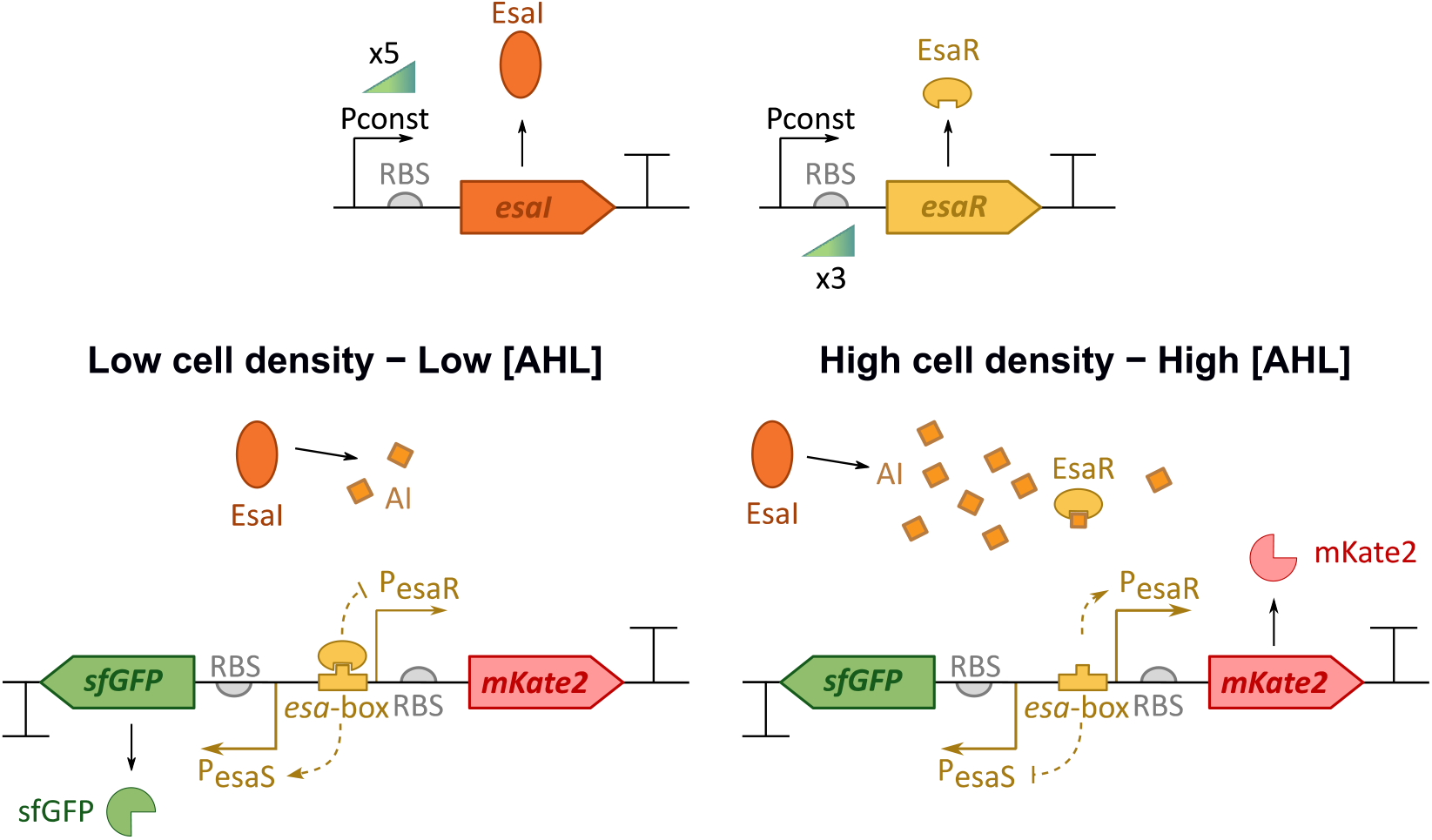
Overview of the EsaI/EsaR quorum sensing system. The transcription factor EsaR binds the promoter region in the absence of 3-oxo-hexanoyl homoserine lactone (3OC6-HSL), its autoinducer (AI), and thereby represses P_esaR_ and activates P_esaS_ which control the expression of the fluorescent reporter proteins mKate2 and sfGFP, respectively. At high cell densities, the AI concentration increases and EsaR gets titrated away from the promoter region. The synthase EsaI and transcription factor EsaR are constitutively expressed (Pconst). For this library five different promoters were combined with *esaI* and three different ribosome binding sites (RBS) with *esaR*, resulting in 15 different strains. Genetic parts are depicted according to SBOL conventions (Hair, 2019; Quinn et al., 2015).

When implementing new genetic parts, synthetic biologists rely on the Design-Build-Test-Learn (DBTL)-cycle to iteratively tune and optimize a system until the desired outcome is obtained. For implementing the EsaI/EsaR system, both the expression level of the synthase and the transcription factor can be varied to tune the dynamics of the system. This results in a large range of possible outcomes that can be obtained from this one system. However, this also implies that the screening of all different combinations to find the desired outcome is labor-intensive. Especially, due to the complex balance between the AHL production by the synthase and the transcription factor level, it is impossible to rationally predict the impact of tuning both parameters simultaneously. This DBTL-cycle can be accelerated with the creation of a predictive mathematical model which aids in unraveling the search space (Dray et al., 2022; Kitano et al., 2023).

Various mechanistic models have been created to capture the dynamics of QS systems. Some models focus on one aspect of the QS system, for example the protein-DNA interactions or the influence of noise on the system (Cox et al., 2003; Mondal and Chaudhury, 2022; Tanouchi et al., 2008), while others capture the full QS network of certain organisms (Dockery and Keener, 2001). Two main distinctions can be made for mathematical models describing QS systems. In general, mechanistic models can be divided in deterministic (Weber and Buceta, 2013) and stochastic models (Cox et al., 2003; Mondal and Chaudhury, 2022; Tanouchi et al., 2008; Weber and Buceta, 2013). For QS systems specifically, a second subdivision can be made between models focusing on the single cell level (Melke et al., 2010; Weber and Buceta, 2013) or the population (Ábrahám et al., 2024; Ward et al., 2001). Most models are not corroborated by biological data. Nevertheless, they still provide meaningful insights in the dynamics of the system and can determine the influence of certain parameters.

In this work, we aim to create a deterministic model that can aid in optimizing the predictability of tuning efforts of the EsaI/EsaR quorum sensing system. Furthermore, the model was fitted to biological data to improve its practical implementation. By varying the expression level of EsaR and EsaI, a library was created that was used for parameter estimation of the model (Figure 1). With the expression level of EsaR and EsaI as inputs, our model aims to predict the response of the P_esaR_ and P_esaS_ promoter.

## 2. Results and discussion

### 2.1 Deterministic model

The developed model comprises eight ordinary differential equations (ODEs) representing the dynamics of the EsaI/EsaR QS system.

The constitutive expression of EsaI is described by Equation 1. The expression strength of the promoter controlling the transcription of *esaI* is varied across the library. The relative strength of this expression is given by the parameter *P* and multiplied with *β*_*Esal*_ to obtain the absolute expression. The term *α*_*Esal*_ is used to take into account that the quantification with the parameter *P* cannot capture the full complexity of EsaI production. Additionally, a term for the degradation and dilution of this protein is included with the degradation rate *k*_*dgrEsal*_.

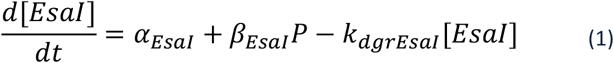

Similarly, the EsaR concentration is captured by Equation 2. Its translation is varied by three different RBS sequences, whose relative strength is approximated by the *R* parameter. Additionally, terms for the association (*k*_1_) and dissociation (*k*_2_) of the EsaR-AHL complex were included.

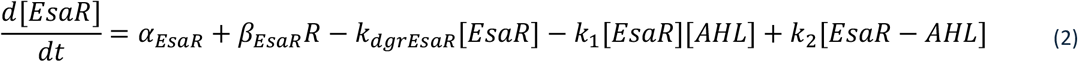

Assuming that the substrate is present in excess, the intracellular AHL production by the EsaI synthase is only dependent on the synthase concentration and the production rate *v*. We assume that the diffusion of the AHL through the cell membrane is not a time-limiting factor. Therefore, the intra- and extracellular AHL-concentration can be considered equal and the molecules are produced by all cells. To achieve this, the production is multiplied by the number of cells at that timepoint (*N*_*c*_(*t*)) and corrected for the ratio of intra-(*V*_*c*_) and extracellular (*V*_*tot*_ − *N*_*c*_(*t*)*V*_*c*_) volume (Equation 3). The full derivation of this correction factor is given in Supplementary Text S1. Additionally, the AHL concentration will be influenced by the association and dissociation of the EsaR-AHL complex.

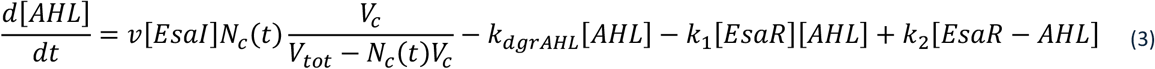

where *N*_*c*_(*t*), the concentration of cells in time is estimated by the Richards function (Equation 4), fitted to the data (Zwietering et al., 1990).

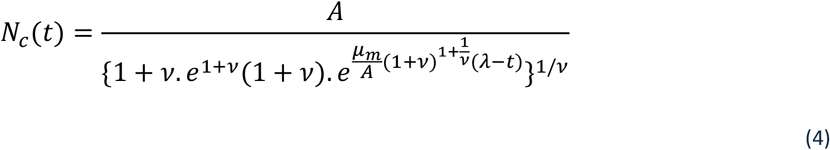

with *A* the initial population density, µ_*m*_ the maximal specific growth rate, λ the lag time and ν a shape parameter.

Equation 5 represents the change in concentration of the EsaR-AHL complex.

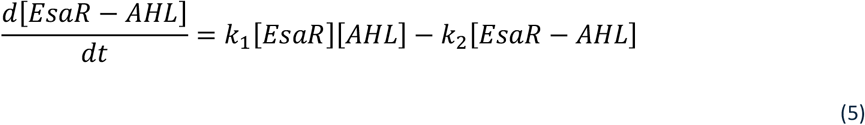

The response of the EsaI/EsaR quorum sensing circuit is monitored by the production of two fluorescent protein reporters: mKate2 for P_esaR_ and sfGFP for P_esaS_. P_esaR_ and P_esaS_ are, respectively, repressed and activated by EsaR, not bound by AHL. Therefore, the promoter activity can be simulated by a Hill function that describes the promoter activity in function of the concentration of the transcription factor (Equation 6 and 7) (Hill, 1913; Santillán, 2008).

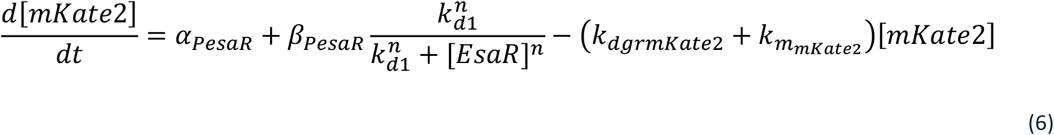

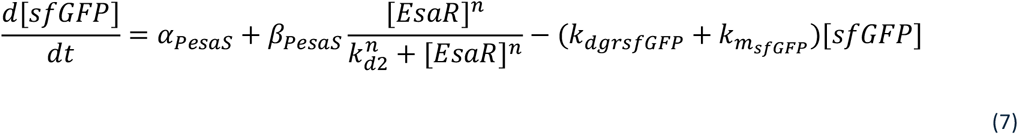

Lastly, the produced protein will mature into its fluorescent variant, described by Equation 8 and 9. The degradation of non-mature and mature fluorescent protein is assumed to be the same. sfGFP was fused to a degradation tag to increase its turnover, this allows to observe the deactivation of P_esaS_.

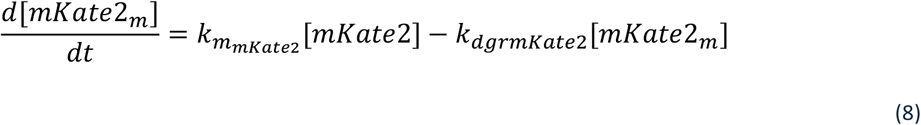

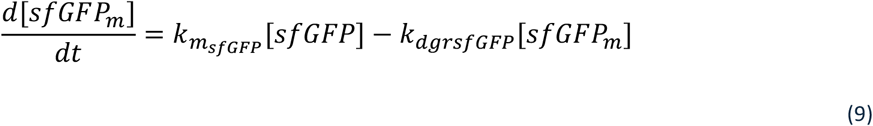

Experimental data was obtained for these final two variables. To allow the comparison, the resulting [*mKate*2_*m*_] and [*GFP*_*m*_] were scaled by multiplication with *s*_*mKate*2_ or *s*_*sfGFP*_, respectively. More information about how these parameters were obtained can be found in the Material and methods.

### 2.2 Fitting the model to experimental data

The main tuning elements in this quorum sensing system are the expression level of the synthase, EsaI, and the transcription factor, EsaR. The synthase level determines how much AHL is produced. The amount of transcription factor is related to how much AHL is needed for the transcription factor to release the promoter. Therefore, it is hard to predict how changing both parameters simultaneously changes the outcome of the system, namely, the timing and final expression level strength. Hence, the purpose of the model created in this work is to improve predictability of the tuning possibilities of the EsaI/EsaR quorum sensing system. By combining the mathematical model with experimental data, this gap can be bridged. For this, a library of 15 variants of the EsaI/EsaR system was created (Figure 1 and Supplementary Figure S1 and S2). Five different promoters, regulating the transcription of *esaI*, were combined with three variants of the RBS controlling the translation of *esaR*. The five promoters were picked from the Anderson promoter collection from the iGEM part registry and selected in the lower range of strengths of that library, since it was shown in earlier research that high levels of EsaI hamper functionality of the system and lead to a significant cellular burden (preprint: De Baets et al., 2025). The three RBS sequences were designed with the Salis RBS calculator with a theoretical strength of 100.18, 529 and 3,115 a.u., referred to as low, medium and high, respectively (Salis, 2011).

### Parameter quantification

To incorporate these expression strengths into the mathematical model, quantification is needed. Even though the theoretical strengths of these regulatory parts are known, the genetic context can lead to major deviations from this prediction (Mutalik et al., 2013). Therefore, fusion proteins of EsaR with sfGFP and of EsaI with sfGFP were created. The fluorescence intensity of these strains was measured as a proxy for the expression level of these proteins. The weakest promoter in the promoter library (Bba_J23117) could not be quantified, because the fluorescence could not be distinguished from the background fluorescence of the wild type strain (Figure 2A). Similarly, protein levels of EsaR-sfGFP whose translation was controlled by the weakest RBS could not be successfully quantified (Figure 2B). Since these parameters could not be accurately estimated, the model for strains containing either this weak promoter or RBS is lacking the *P* or *R* parameter and cannot be fit. Therefore, these strains were removed from the library, resulting in a final library size of eight strains, compared to the originally planned fifteen.

**Figure 2.**
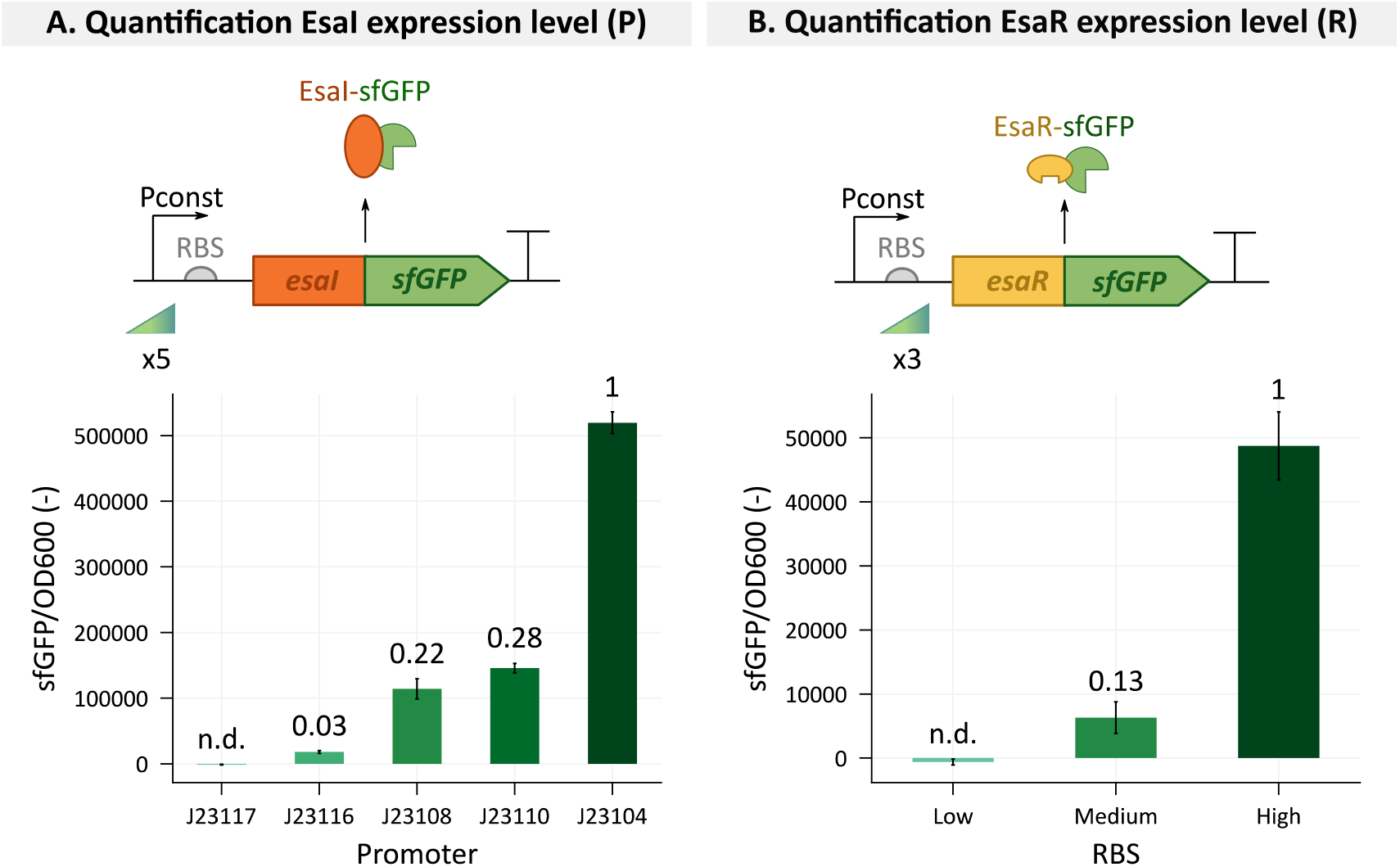
Quantification of the expression strengths of the promoters and ribosome binding site (RBS) sequences controlling the expression of *esaI* (**A**.) and *esaR* (**B**.), respectively. To achieve this, both proteins were fused with sfGFP and the resulting green fluorescence was measured as a proxy for protein levels, reflecting the expression strength. The relative expression strengths, corresponding to the P and R-values, were added on top of each bar. Fluorescent values were obtained in the stationary phase and normalized for cell growth determined by optical density at 600 nm (OD600). Bars represent the mean and error bars the standard error of the mean based on three biological replicates. Genetic parts are depicted according to SBOL conventions (Hair, 2019; Quinn et al., 2015).

Besides the expression levels of EsaI and EsaR, the model also requires the number of cells in function of time as an input. Since the growth function is independent of the rest of the model, the parameters in the Richards growth curve were estimated separately and the resulting parameters were fixed as input for the QS model. By taking the growth fitting out of the model, we reduce the model complexity. The obtained fits and the corresponding parameter values are collected in Supplementary Figures S5 and S6 and Supplementary Table S1. The resulting parameters were then provided to the model to replicate the growth of each specific strain. Since this curve was fitted to optical density values and the model requires the actual number of cells, a calibration curve derived from De Wannemaeker *et al*. (2023) (De Wannemaeker et al., 2023), was used to transform the optical density values to cell quantity.

### Model fit

The remaining eight strains were divided over a training (six strains) and test (two strains) set. The training set was used for parameter estimation by minimizing the residuals between the experimental data and the model output. In a first attempt to fit the model to this training set, 17 parameters were fixed beforehand based on values derived from literature (Table 1). The six remaining parameters, *i*.*e*., the production rates of EsaI and EsaR and the Hill constants of P_esaR_ and P_esaS_, were allowed to be varied across a biologically relevant range.

**Table 1.** Overview of the parameter values of the model describing the EsaI/EsaR quorum sensing system.

| Parameter | Meaning | Value | Unit | Reference |
| --- | --- | --- | --- | --- |
| $\alpha_{\text{EsaI}}$ | Basal *EsaI* expression rate | 92.77 | nM/h | Fitted |
| $\beta_{\text{EsaI}}$ | Basal *EsaI* expression rate | 647.37 | nM/h | Fitted |
| $\alpha_{\text{EsaR}}$ | Basal *EsaR* expression rate | 15 | nM/h | Fitted & fixed |
| $\beta_{\text{EsaR}}$ | Basal *EsaR* expression rate | 18 | nM/h | Fitted & fixed |
| $\alpha_{P_{\text{EsaR}}}$ | Basal expression $P_{\text{EsaR}}$ | 1 | nM/h | Fixed |
| $\beta_{P_{\text{EsaR}}}$ | Maximal promoter activity $P_{\text{EsaR}}$ | 1000 | nM/h | Fixed |
| $\alpha_{P_{\text{EsaS}}}$ | Basal expression $P_{\text{EsaS}}$ | 1 | nM/h | Fixed |
| $\beta_{P_{\text{EsaS}}}$ | Maximal promoter activity $P_{\text{EsaS}}$ | 1000 | nM/h | Fixed |
| $k_1$ | Association rate *EsaR*-AHL | 0.545 | 1/(nM.h) | (Fagerlind et al., 2005) |
| $k_2$ | Dissociation rate *EsaR*-AHL | 0.001 | 1/h | Assumption based on (Saeidi et al., 2013) |
| $v$ | AHL production rate of Esal | 10 | nM/h | Assumption based on (Schaefer et al., 1996; Weber and Buceta, 2013) |
| $k_{d1}$ | Hill constant $P_{\text{esaR}}$ | 0.89 | nM | Fitted |
| $k_{d2}$ | Hill constant $P_{\text{esaS}}$ | 328.13 | nM | Fitted |
| $n$ | Hill coefficient Esar | 2 | / | (preprint: De Baets et al., 2025; Schu et al., 2009) |
| $k_{dgr\text{Esal}}$ | Degradation rate Esal | 0.05 | 1/h | (Boada et al., 2020) |
| $k_{dgr\text{Esar}}$ | Degradation rate Esar | 0.05 | 1/h | (Boada et al., 2020) |
| $k_{dgr\text{Esar-AHL}}$ | Degradation rate Esar-AHL | 0.01 | 1/h | Assumption |
| $k_{dgr\text{AHL}}$ | Degradation rate of AHL | 0.0044 | 1/h | Assumption based on (Fekete et al., 2010) |
| $k_{dgr\text{mKate2}}$ | Degradation rate mKate2 | 0.5 | 1/h | Assumption |
| $k_{dgr\text{sfGFP}}$ | Degradation rate sfGFP | 0.8 | 1/h | Assumption based on (Claussen et al., 2013) |
| $k_{m\text{mKate2}}$ | Maturation rate mKate2 | 3 | 1/h | Assumption based on (Balleza et al., 2018; Shcherbo et al., 2009) |
| $k_{m\text{sfGFP}}$ | Maturation rate sfGFP | 1 | 1/h | (Sun et al., 2024) |
| $V_c$ | Cell volume *E. coli* | 1e-9 | μL | (Milo, 2013) |
| $V_{tot}$ | Reaction volume | 150 | μL | / |
| $S_{m\text{Kate2}}$ | mKate2 scaling factor | 0.00041703 | / | Calculated |
| $S_{m\text{sfGFP}}$ | sfGFP scaling factor | 0.00142421 | / | Calculated |

During the optimization process, it could be observed that the six initial parameter values had a major influence on the resulting parameter estimates and model quality. This indicates that this complex six-dimensional parameter space contains a lot of local minima and the tested minimization methods were not successful in avoiding these to find the global minimum. Hence, despite the numerous efforts by varying the initial parameter estimates and testing multiple minimization methods, the obtained fitted model was not able to accurately predict the behavior of all eight strains in the training set (Supplementary Figure S5). Especially the highest expression level of the synthase with promoter Bba_J23104 could not be accurately fit without compromising the fits of the other strains. The explanation can be found in the strength of this promoter compared to the rest of the library, where the Bba_J23104 promoter already leads to 3.5 times more EsaI production than the Bba_J23110 promoter. Additionally, in these strains, almost no P_esaS_ activity can be observed. Therefore, we reason that the EsaI level already leads to threshold AHL-concentrations in the beginning of the growth process. Furthermore, it is assumed that once the EsaI expression level surpasses a certain level, the same saturated behavior would be observed. Hence, once the *P*-parameter is higher than a certain value, the observed output would be the same. Therefore, the data provided by these strains would not contribute to the quality of the fit. However, by removing these strains from the library, only six different strains remain. Since the final library size got drastically reduced, the remaining strains were not split up over a training and test set anymore. We hypothesize that fitting the model to the six different strains with different characteristics that cover the range of possible outcomes, minimizes the risk of overfitting.

The parameters were now estimated by fitting the model to the six remaining strains (Supplementary Figure S6, Supplementary Table S2). However, correlations between most of the parameters can be found (Table 2). Especially β_*EsaR*_ appears to be correlated with all parameters besides *k*_*d*1_ and *k*_*d*2_. From a biological perspective, this correlation is not surprising. Theoretically, for each level of the transcription factor, there exists an EsaI level that results in the necessary amount of AHL to obtain the observed results. Nevertheless, not all expression levels are possible from a biological perspective. These correlations might result in parameters that are structurally unidentifiable. This implies that the structure of the model does not allow for the estimation of the best set of parameters to explain the data. This does not per se mean that the model is not valuable, but it might pose issues when interested in variables that are currently not observed (such as the AHL-concentration) (Muñoz-Tamayo et al., 2018).

**Table 2.**
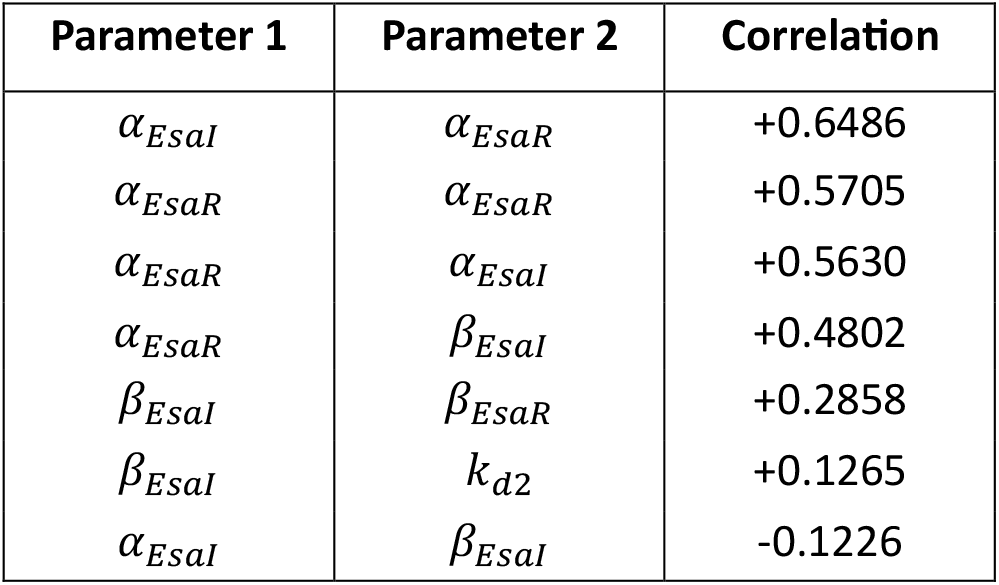
Overview of the correlations found between the estimated parameters. Correlations below 0.1 are not given.

The identifiability issues are confirmed by the Markov Chain Monte Carlo (MCMC) analysis performed on the parameters to find their posterior distributions (Figure 3). The traces of the corresponding walkers are given in Supplementary Figure S7. The analysis did not succeed in finding symmetric, unimodal parameter distributions because of the strong correlations between all the parameters, as can be seen in the two-dimensional density plots. The identifiability issue can be resolved by changing the structure of the model, *e*.*g*., model reduction obtained by fixing parameters, but it has the risk of decreasing the accuracy of the model. As mentioned earlier, especially the expression level of EsaR appears to be correlated to the other parameters. Additionally, the estimated expression strength of EsaR in the current fit, results in EsaR levels that correspond to typical transcription factor levels found in nature, namely 1-1000 nM (Ishihama et al., 2008). Therefore, both α_*EsaR*_ and β_*EsaR*_ were fixed to 15 and 18 nM/h, respectively.

**Figure 3.**
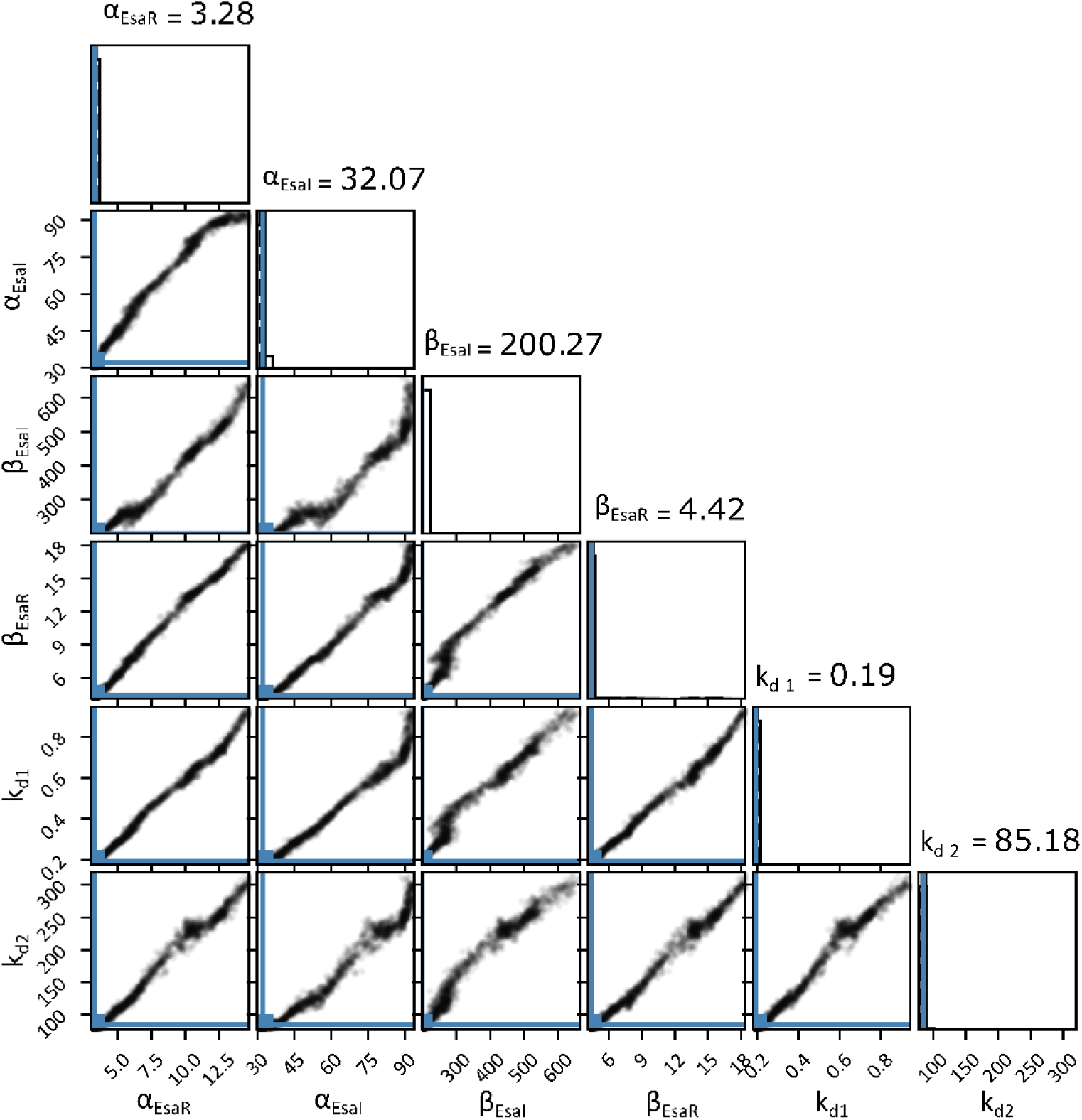
Posterior distributions of the estimated parameters obtained via Markov Chain Monte Carlo analysis. Above the distribution, the parameter estimate is given.

With these two parameters now fixed, the residuals were again minimized. The resulting parameters were analyzed with MCMC to identify the optimal parameter estimates and their posterior distributions (Figure 4). The traces of the corresponding walkers are given in Supplementary Figure S8. For all four parameters, a unimodal and symmetric distribution was obtained which allowed for the calculation of credible intervals. This indicates that the model is now identifiable. The analysis revealed that there are still correlations between the parameters: β_*Esal*_ and α_*Esal*_ show a negative correlation (quantified at −0.6278) and lower correlations were found between β_*Esal*_ and *k*_*d*2_ (+0.1939) and between α_*Esal*_ and *k*_*d*1_ (−0.1359). The negative correlation between β_*Esal*_ and α_*Esal*_ is not that surprising, because together these two parameters determine the EsaI expression level. Hence, if one increases, the other parameter has to decrease to keep the total EsaI level in the same range. The final estimated parameter values and their corresponding credible interval are given in Figure 4.

**Figure 4.**
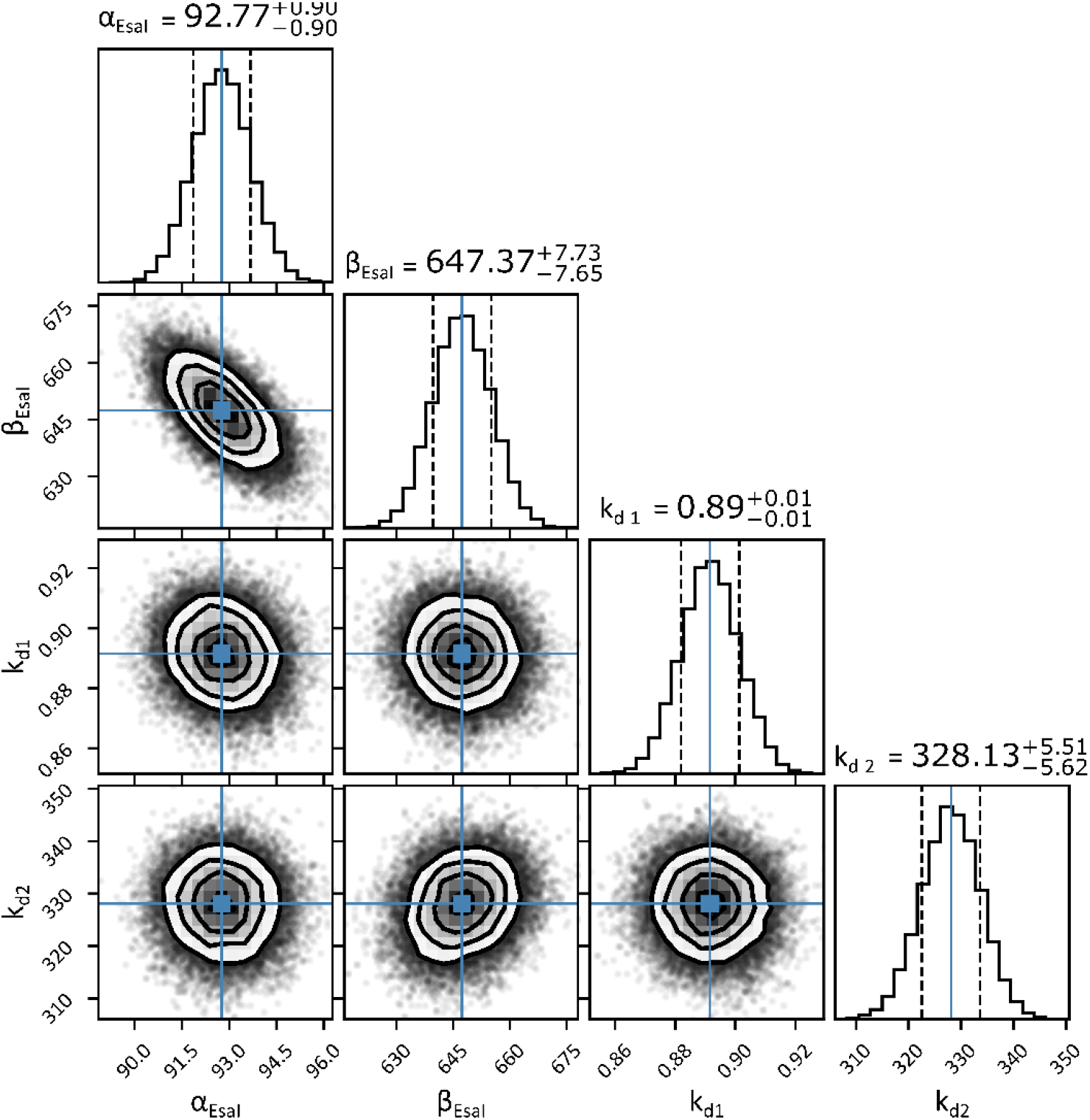
Posterior distributions of the estimated parameters obtained via Markov Chain Monte Carlo analysis. Above the distribution, the parameter estimate and corresponding 68% credible interval are given, also highlighted by the vertical blue and dashed black lines, respectively. On the density plots, the (0.5, 1, 1.5, 2)-sigma equivalent contours are drawn, *i*.*e*., containing 11.8%, 39.3%, 67.5% and 86.4% of the samples.

The resulting fit of the strains in the library is visualized in Figure 5. This model succeeds in following both the mKate2 and sfGFP dynamics and can approximate the switching time of all strains. Only for the strains with promoter Bba_J23116 and the high strength RBS, the mKate2 upregulation is predicted to be later than in the observed data. This model also captures differences in the height of the sfGFP peaks. Nevertheless, the actual height of the peak cannot be accurately predicted for all strains. Similarly, the final mKate2 levels of the different strains cannot be accurately predicted. At this time point, the strains have reached the stationary growth phase, which besides the halted growth, also comes with large metabolic changes which influence the promoter activity (Rothschild et al., 2014). During this growth stage, there is an 80% reduction in protein synthesis, mainly related to the lower activity of the housekeeping sigma factor (Reeve et al., 1984). Consequently, quorum sensing strains which switch closer to the stationary phase, will not be capable of accumulating as much mKate2 as strains with an early switch before the P_esaR_ activity is reduced. Nevertheless, the effect of the stationary phase is currently not included in the deterministic model. One option to incorporate this in future models is by including a time-dependent repression term. As such, this term, which would decrease with increasing cell density, could be multiplied with the promoter activity to reduce the promoter activity in the stationary phase (Klumpp et al., 2009).

**Figure 5.**
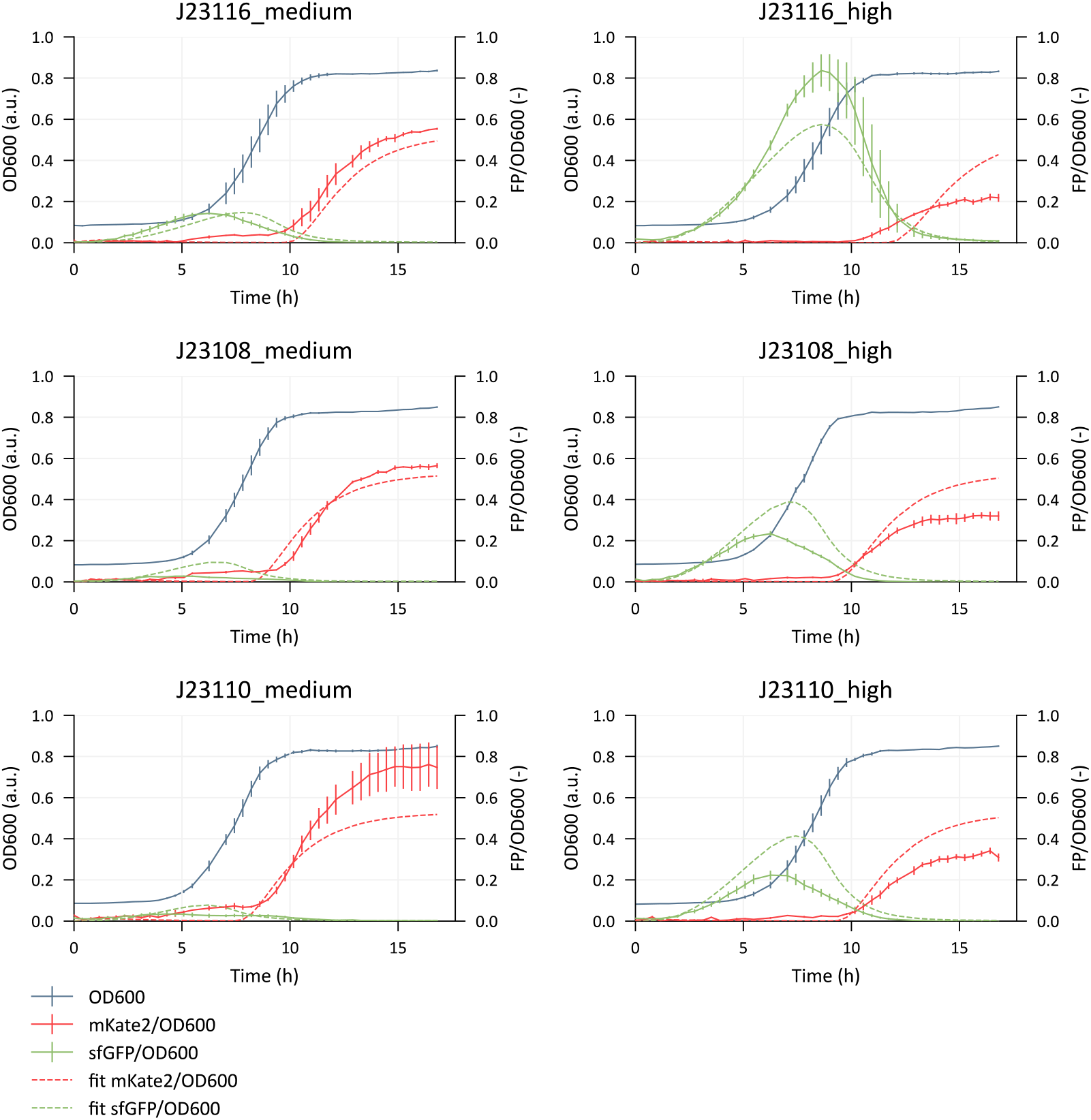
Model fit to the six strains of the EsaI/EsaR library. The dashed lines depict the prediction made by the model. Fluorescent values are normalized for cell growth determined by optical density at 600 nm (OD600). Error bars represent the standard error for three biological replicates. FP = fluorescent protein, referring to either mKate2 or sfGFP.

### Global sensitivity analysis

To gain more insight into the influence of each estimated parameter on the output of the EsaI/EsaR system, a global Sobol sensitivity analysis was performed (Sobol, 2001). This analysis was used to evaluate how variations in each parameter within its credible interval contributed to the variability of the output variables, either mKate2 or sfGFP. The results are summarized in Figure 6. This analysis was performed on each strain separately as parameter sensitivity could depend on the *P* and *R* values.

**Figure 6.**
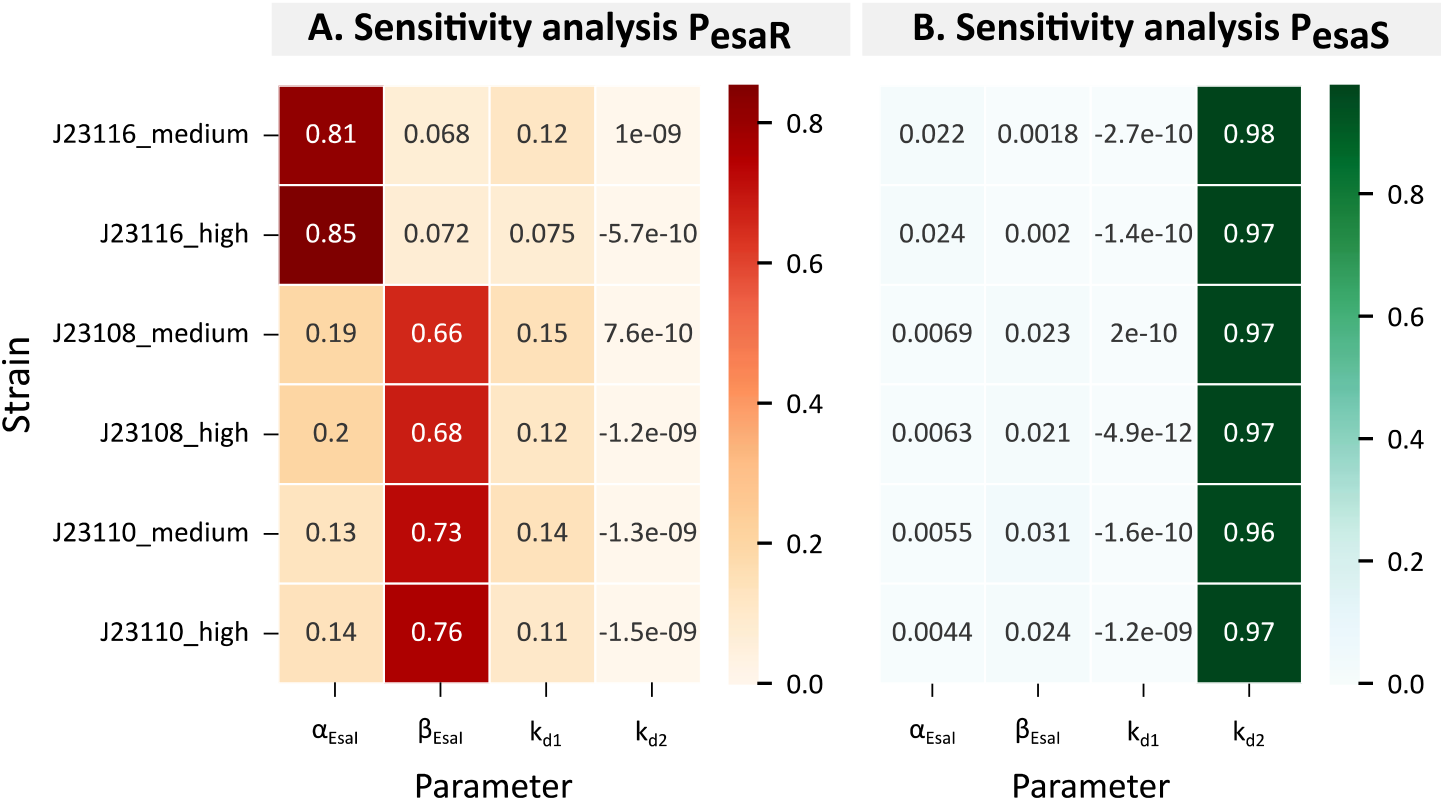
Heatmap of the first order Sobol indices of the global sensitivity analysis of P_esaR_ (**A**.) and P_esaS_ (**B**.). Rows represent the six different strains used in the model fit, columns represent the estimated parameters.

The P_esaR_ activity appears to be most sensitive to the synthase expression level, reflected by α_*Esal*_ and β_*Esal*_. This can be explained by the influence that the synthase level has on the timing of the switch and the upregulation of this promoter. Remarkably, the strains with the Bba_J23116 promoter are more sensitive to changes in the α_*Esal*_-value compared to the β_*Esal*_ parameter. This can be explained by the low *P*-value of 0.035. Hence, the resulting *P* ∗ β_*Esal*_-value is negligible compared to the α_*Esal*_-value. Besides the EsaI expression level, the promoter activity is also, although to a lower extent, sensitive to changes in the *k*_*d*1_-value. From these results can be concluded that for accurate predictions of the P_esaR_ promoter activity, an accurate estimation of the synthase level is of importance, because this will contribute the most to the output. Nevertheless, we also experienced how difficult the estimation of this parameter is due to its correlations to other parameters.

Variations in the Hill constant of the P_esaS_ promoter, *k*_*d*2_, appear to have the most influence on the output of this promoter. High sensitivity to *k*_*d*2_ indicates that the affinity of the transcription factor for its transcription factor binding site (TFBS), located in the promoter region, can be an important tuning parameter for this QS system. This finding is supported by the work of Shong *et al*. (2013) who added an additional TFBS at different locations within the promoter region, resulting in a large influence on the response of P_esaR_ (Shong and Collins, 2013). The influence of variation in the synthase expression level on P_esaS_ activity is shown to be minimal. However, this does not mean that the synthase expression level does not influence the promoter output, because this behavior is captured by the quantified *P* parameter. But, within the credible intervals of the estimated parameters, especially *k*_*d*2_, is of importance.

It is of importance to note that this sensitivity analysis is relevant for this specific model. If other assumptions were made or different parameter values were used, the sensitivity analysis could result in different observations.

### Parameter values discussion

The obtained parameter estimates do not necessarily correspond to the true biological value for various reasons. First, models are a simplification of the true biological process. As such, processes such as transcription and translation were not described by separate differential equations in this model. Additionally, EsaR binds the promoter region as a dimer, which was also not incorporated in the model. Secondly, the model is based on differential equations and cannot account for the complex processes occurring in the cell that might influence the final output of the QS system. This can relate to stress and competition for resources, but also the effect of the growth phase on promoter activity. These factors were all not explicitly incorporated into the model, yet the model succeeded in reasonable predictions of the output of the QS system. This implies that these factors were compensated for in different ways. Thirdly, the values of the fixed parameters and the initial parameter estimates of the optimized parameters, all derived from literature, had a big influence on the obtained model fit. However, the parameter values found in literature show large variations across research papers. Hence, the source used for these initial parameters will impact the final parameter estimates. In conclusion, interpreting the obtained parameter values must be done with care. Therefore, it is important to remember the famous expression by statistician George Box: ‘All models are wrong, but some are useful’ (Box, 1976).

Nevertheless, the obtained parameter estimates can contain valuable information about the QS system. Interestingly, the *k*_*d*1_ and *k*_*d*2_ are estimated at completely different values. Repression is generally a stronger process than activation, which could be an explanation for the lower *k*_*d*_ value for the repression of P_esaR_. However, theoretically, the two *k*_*d*_-values should be the same as they represent binding of EsaR to the same promoter region. The difference in the obtained values indicates that these values overcompensate for effects that cannot be captured by the simplistic model. Furthermore, both the *k*_*d*1_- and *k*_*d*2_-value are estimated near their lower and upper bound, respectively. It was observed that when looser bounds were used, the minimization process got stuck at a local minimum with inaccurate fittings. Therefore, the stricter bounds were retained. This again confirms that the obtained parameter values need to be interpreted with care and that the simplicity of the model is compensated by parameter estimates that are not biologically accurate.

It can also be observed that the estimated α_*EsaR*_ and β_*EsaR*_ are in the same range. This indicates that according to the model, the differences in EsaR levels between the strains are not as high as measured during the parameter quantification. We reason that the truth is probably somewhere in between. As mentioned earlier, this model is only an approximation which does not consider many of the biological processes influencing protein levels and the dynamics of the system. Moreover, the method we used to quantify the *P* and *R* values does not involve direct measurements and may be affected by various factors and the detection limit. For α_*Esal*_ and β_*Esal*_ the difference appears bigger, however, *P* is estimated at 0.28 for strains with promoter Bba_J23110, the strongest promoter after the removal of Bba_J23104 from the library. Hence, the final expression strength of EsaI is explained for a maximum of 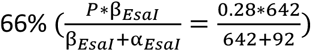 by the quantified promoter strength.

Most of the parameters in this model were fixed on values based on literature. However, large discrepancies exist between the reported values, which can again be explained by the relevance of a certain parameter value within a certain model and experimental set-up. The parameters representing the expression strength of the QS-regulated promoters were kept constant, because the fluorescent values were normalized anyway. Furthermore, the relation between these parameters and the observable output, the fluorescence, is more straightforward and does not benefit from being incorporated into this model.

### 2.3 Assessing the performance of the model

Theoretically, the obtained model should be capable of both predicting and forward engineering the EsaI/EsaR system. This means that starting from a quantified relative expression level of EsaI and EsaR, the model would predict the P_esaR_ and P_esaS_ response in time. Secondly, given a certain response in time, the model could predict the relative expression level of EsaR and EsaI.

To get a better view of the influence of the EsaI and EsaR expression level on the response of the system, 25 theoretical combinations were assessed. Figure 7 gives a full overview of the theoretical possibilities of the EsaI/EsaR system and the corresponding biological design parameters (*P* and *R*). The height of the sfGFP peak and the induction timing of mKate2 vary the most across the 25 variants. Again, differences in mKate2 levels in the stationary phase cannot be observed across the strains, but as discussed earlier this is related to a flaw in the mathematical model.

**Figure 7.**
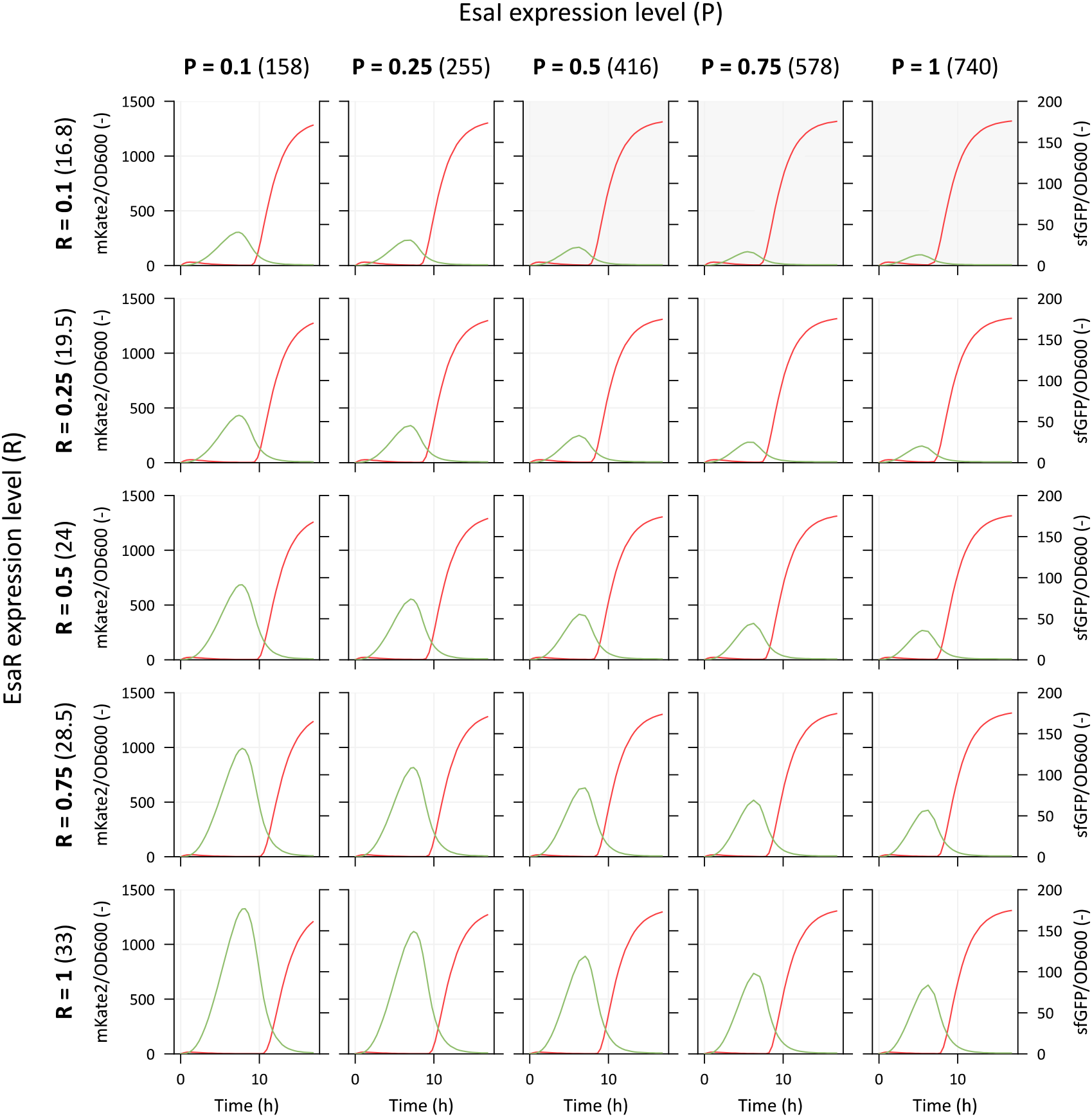
Overview of the P_esaR_ and P_esaS_ response, regulating mKate2 and sfGFP expression, respectively, for different expression levels of the synthase (P) and the transcription factor (R). The value in brackets represents the total expression level (α + (P or R)^*^β). The subplot in gray represents strains with sfGFP peaks that are most likely not detectable. OD600 = optical density at 600 nm.

Remarkably, for a fixed *P*-value, the output differs more with varying *R*-values compared to when *R* is kept constant and *P* varied. Additionally, the corresponding expression level (α + (*P or R*) ∗ β) covers a bigger range of values for EsaI expression compared to EsaR. This further accentuates the importance of the EsaR expression level on the response. A small difference in its expression strength has the largest influence on the output. In contrast, varying the expression level of the synthase protein is until now the most used tuning technique for QS systems (Dinh et al., 2020; Dinh and Prather, 2019; Gupta et al., 2017; He et al., 2017). It can also be observed that a similar response can be obtained by different combinations of *P* and *R, e*.*g*., the system with *P* = 0.5 and *R* = 0.5 has a similar response as the system with *P* = 1 and *R* = 0.75. This also indicates that it would not be possible to apply our model for forward engineering, since there are multiple possible solutions for the model to result in given response.

Not all the combinations in this grid (Figure 7) might be biologically feasible. For example, for the strain with promoter Bba_J23110 (*P* = 0.28) and medium strength RBS (*R* = 0.13), the sfGFP peak can almost not be observed and distinguished from the background fluorescence (Figure 5). Hence, strains with *P*- and *R*-values higher than these (colored gray on Figure 7) will most likely not have observable P_esaS_ activity. Furthermore, other combinations that are not indicated on the figure might have the same issues.

To fully assess the performance of the final model, assessing its predictive performance needs to be assessed for new strains that lie in the range used for training the model (*P* = 0.035 to 0.28 and *R* = 0.13 to 1) and strains that lie outside of that range.

## 3. Conclusions

The EsaI/EsaR quorum sensing system is an interesting starting point for synthetic biology applications, such as biosynthesis pathway regulation. Nevertheless, tuning of quorum sensing systems, and genetic circuits in general, remains a cumbersome undertaking. This can be related to multiple factors. Firstly, the lack of standardized and characterized parts remains a bottleneck in the synthetic biology toolbox, thereby hampering the predictability of the resulting engineered strain. Secondly, in QS systems, the intricate balance between transcription factor and synthase levels remains poorly understood, often restricting scientists to adjusting only one of the two parameters. Consequently, this narrows down the range of possible outcomes. By implementing a mathematical model, we aim to unravel this balance and improve the predictability of tuning these circuits on multiple levels simultaneously. This deterministic model was fit to experimental data obtained from six strains with different expression levels of transcription factor and synthase. Nevertheless, the complex relation between the synthase and transcription factor levels complicated this fitting process due to their high correlation. Yet, a model was obtained that could predict the induction timing and differences in P_esaR/esaS_ expression levels. The resulting model experienced difficulties with accurately estimating the actual promoter activity of both promoters. This issue could be attributed to the influence of the stationary phase that was not included in the current model. Nonetheless, the model can still be a valuable tool in predicting the general response of the system and differences between varying strains. Interestingly, we observed that the EsaR expression level appeared to be an easier tuning parameter than the synthase expression level. Small variations in its expression could already change the output.

In conclusion, a mathematical model was created that aims to improve the predictability of tuning strategies of the EsaI/EsaR quorum sensing system. Up until now, most QS models focused on unraveling more about the dynamics and details of these systems. In contrast, the model created here really aims at bridging the gap between theoretical tuning strategies and the actual *in vivo* result. However, its applicability is now still limited to the genetic regulatory parts used in this research and can be expanded when new promoters or RBS-sequences are characterized relative to the parts in this research. Nevertheless, we envision higher applicability in a future world with standardized and characterized genetic parts.

## 4. Material and methods

### 4.1 Strains and media

Enzymes and related products are purchased from New England Biolabs (County Road, Ipswich, MA, USA), other chemicals from Sigma-Aldrich (Brusselsesteenweg, Overijse, Belgium) unless stated differently. Protocols as described by the vendors were applied unless mentioned otherwise.

Newly assembled plasmids were introduced in One Shot® Top10 Chemically Competent *E. coli* cells (Invitrogen, Carlsbad, California, USA). The fluorescence experiments were performed in *E. coli* K12 MG1655 cells (mutant version of ATCC 47076, additional deletions resulting in Δ*ynaJ* Δ*uspE* Δ*fnr* Δ*cct* Δ*abgT* Δ*abgB* Δ*abgA* Δ*abgR* Δ*smrA* Δ*ydaM* Δ*ydaN* Δ*dbpA* Δ*ttcA* Δ*intR* Δ*recT* Δ*recE*).

Lysogenic Broth (LB) was used for growth during the cloning process and for preculture plates for microtiter plate experiments. This medium is composed of 10 g/L Tryptone (BioKar Diagnostics, Allonne, France), 5 g/L Yeast Extract (Becton Dickinson, Erembodegem-Dorp, Erembodegem, Belgium) and 5 g/L NaCl. 12 g/L agar (BioKar Diagnostics, Allonne, France) is added to LB to make LB-agar plates. Kanamycin was added when necessary in a 1000 times dilution of the filter sterilized stock solution resulting in a final concentration of 50 µg/mL. Cultures were grown at 30°C at 200 rpm (LS-X (5 cm orbit), Adolf Kühner AG, Switzerland). Super Optimal Broth (SOB medium) was used for growing the overnight *E. coli* cultures to make them chemocompetent and for the resuscitation step after heat shock transformation. The medium consists of 20 g/L Tryptone (BioKar Diagnostics, Allonne, France), 5 g/L Yeast Extract (Becton Dickinson, Erembodegem-Dorp, Erembodegem, Belgium), 0.5 g/L NaCl, 2.5 mM KCl and 10 mM MgCl_2_. Experiments were performed in MOPS EZ Rich Defined Medium (M2105) (Teknova, Hollister, CA, USA) with 0.2% glucose as carbon source. The medium was prepared according to the protocol provided by the vendor.

Phosphate buffered saline (PBS) (P5368) from Sigma-Aldrich (Brusselsesteenweg, Overijse, Belgium) was used to wash the precultures to remove all autoinducers.

### 4.2 Plasmid construction

An overview of all the used plasmids is given in Table 3. An overview of the DNA-sequence of all regulatory parts and genes is given in Supplementary Table S3 and S4.

**Table 3.**
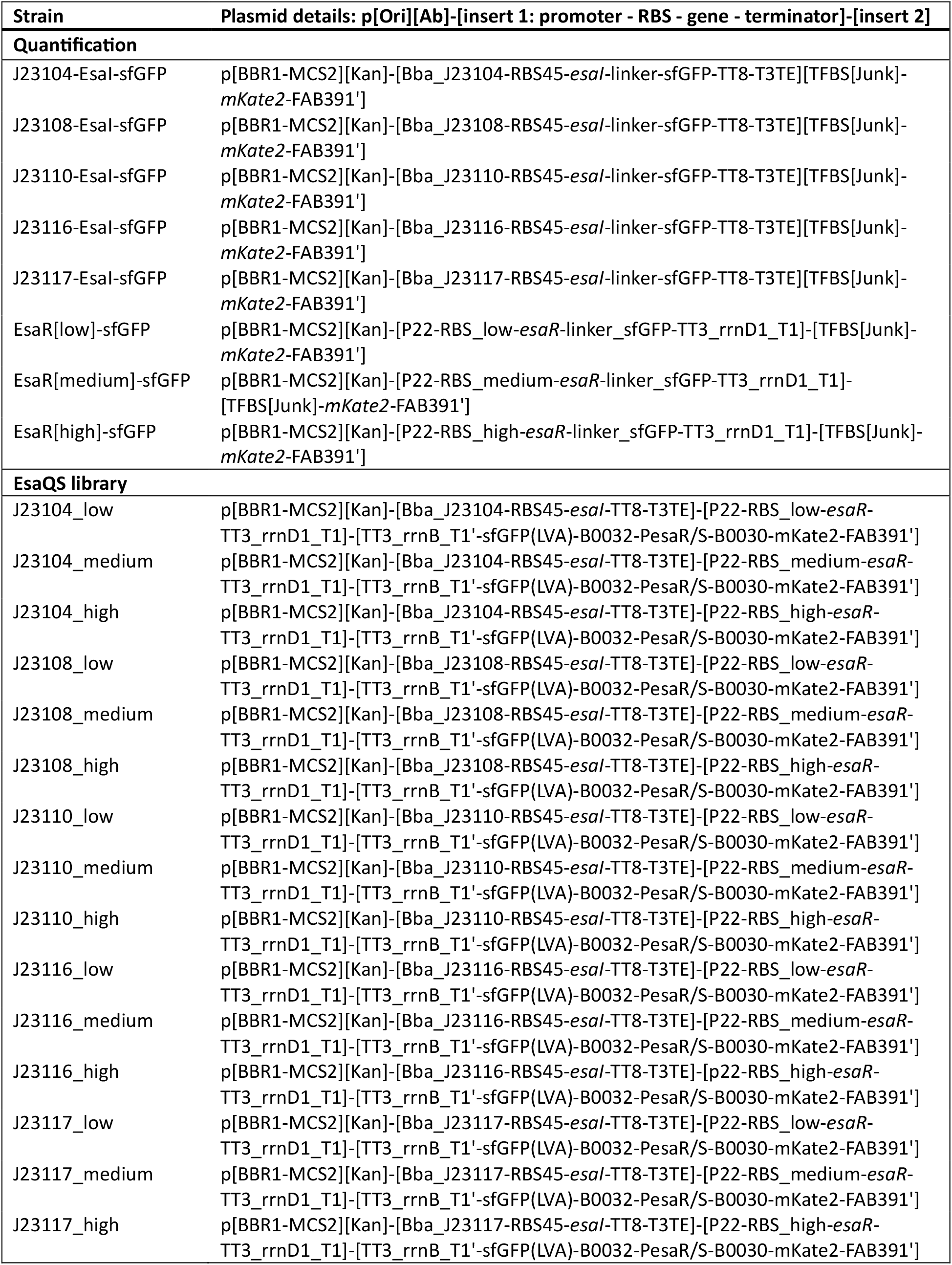
Overview of the plasmids used in this research. All plasmids are made in the MoBioS platform as also described by De Baets *et al*. (2025) (preprint: De Baets et al., 2025; Demeester et al., 2023).

All plasmids were created in the MoBioS backbone with PaqCI Golden Gate as described in Demeester *et al*. (2023) and De Baets *et al*. (2025) (preprint: De Baets et al., 2025; Demeester et al., 2023; Engler et al., 2008).

For the construction of the transcription factor-sfGFP fusions, the transcription factor and a non-functional promoter region were added as inserts for Golden Gate with the Type II restriction enzyme PaqCI (New England Biolabs, Inc., USA) and the MoBioS platform as acceptor plasmid. Afterwards, sfGFP with a glycin-rich linker (GGSGGGSG) was attached to the C-terminal of the transcription factor using Circular Polymerase Extension Cloning (CPEC) with Q5 DNA polymerase (New England Biolabs, Ipswich, MA, USA) (Quan and Tian, 2009).

The EsaI-sfGFP fusion was created starting from the MoBioS-EsaI plasmid from De Baets *et al*. (2025) (preprint: De Baets et al., 2025). Again, sfGFP with a glycin-rich linker (GGSGGGSG) was attached to the C-terminal of this protein via CPEC. Afterwards, the promoter was interchanged with the four new promoters via CPEC. Similarly, the promoter was replaced in the strain without the sfGFP fusion. Next, the EsaR (with the three different RBS sequences) and the promoter region were introduced as linear parts via Golden Gate with the Type II restriction enzyme PaqCI (New England Biolabs, Inc., USA) as described by Demeester *et al*. (2023) (Demeester et al., 2023).

All polymerase chain reactions (PCR) performed were done using PrimeStar HS (Takara, Westburg, Leusden, The Netherlands). A list of all oligonucleotides (IDT, Leuven, Belgium) used during this project is given in Supplementary Table S5.

One Shot® Top10 Chemically Competent *E. coli* cells (Invitrogen, Carlsbad, California, USA) were transformed with the constructed plasmids. Successful transformation was checked by colony PCR. The plasmids of overnight growth cultures of positive colonies were prepped using the QIAprep Spin Miniprep Kit (Qiagen, Venlo, The Netherlands) and sent for Sanger sequencing to Macrogen (Macrogen Inc., Amsterdam, The Netherlands). Finished plasmids were then transformed into *E. coli* K12 MG1655 for the production experiments.

Cells were made chemocompetent with the *Mix & Go! E. coli* Transformation Kit (T3002, Zymo Research, ordered via BaseClear B. V., Leiden, The Netherlands). After adding DNA to pre-chilled cells, the cells were heat-shocked for 45 seconds at 42°C and then placed on ice for 2 minutes, before adding 900 µL SOB for regeneration. After 1-3 hours of resuscitation, the cells were plates on LB-agar plates containing the required antibiotics.

### 4.3 *In vivo* fluorescence experiments

Precultures were inoculated in a transparent flat-bottomed 96-well plate (Greiner Bio-One) filled with 150 µL LB medium supplemented with kanamycin when needed. These preculture plates were incubated at 30 °C for 16 to 18 hours while shaking at 800 rpm using the Digital Microplate Shaker (Thermo Fisher Scientific). Afterwards, the precultures were washed to remove all autoinducers that were produced during the overnight growth. The preculture plates were centrifuged at 4000 rpm for 30 minutes with the Rotanta 46 RSC centrifuge (Hettich Benelux, Geldermalsen, The Netherlands) at 4°C. Supernatant was removed and the pellets were resuspended in 150 µL PBS-solution. The same centrifugation and resuspension step was repeated. Afterwards a 300 times dilution of the washed precultures plates was made into a black flat-bottomed 96-well plate (Greiner Bio-One).

The strains were analyzed in our integrated robotic system. Cultures were incubated in the Inheco Incubator Shaker MP (integrated in the explorer G3 workstation) at 30°C and 800 rpm. Optical density at 600 nm (OD600) and fluorescence measurements were taken every 20 minutes in the PerkinElmer Ensight multimode plate reader (integrated in the explorer G3 workstation). The used excitation and emission wavelengths for mKate2 are 588 and 633 nm, respectively. For sfGFP 480 nm and 510 nm were used for excitation and emission, respectively. The lid of the microtiter plates was coated with a Triton X-100 solution to reduce condensation. The Triton X-100 solution was made by mixing 20 mL ethanol with 80 mL sterile water and adding 50 µL Triton X-100. This solution was poured onto the lid and incubated for one minute. The solution was then discarded and the lid was fully dried in the fume hood before using it.

### 4.4 Data processing

The full code for data processing, model fitting, analysis and visualization is provided at https://github.com/MEMO-group/QuorumSensing_model_JDB.git under an open-source license.

The data was analyzed in Python (version 3.12) using Jupyter Notebooks with the pandas package. The obtained fluorescence values (*Fluo*) were normalized for optical density (*OD*_600_) per time point with Equation 10. Background fluorescence was considered by measuring the optical density (*OD*_600,*WT*_) and fluorescence (*Fluo*_*WT*_) of the wild type strain *E. coli* K12 MG1655 in MOPS EZ Rich defined medium. The corrected and normalized fluorescence was then averaged over the three repeats and the standard error was obtained.

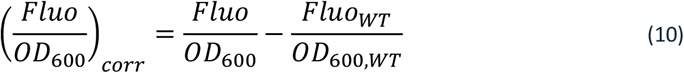

Next, the fluorescence values of the strains in the library were normalized to be in the range between zero and one. For this, the normalized fluorescent values were divided by the maximum normalized fluorescence of the highest producing strain.

For fitting the Richards growth curve to the optical density values, these values were first corrected for background density of the medium.

### 4.5 Model fitting and analysis

The Richards growth curve was fitted to the corrected optical density with the curve_fit package in Python (version 3.12) for each biological replicate separately. Next, the geometrical mean of each parameter estimated was calculated and used as an input for the model to determine the number of cells. The conversion from cell density to cell numbers was done with the calibration curve described by De Wannemaeker *et al*. (2023) specifically for the used plate reader (De Wannemaeker et al., 2023).

Before calculating the residuals between the experimental and model outcome, the model outcome was scaled to be in the same range as the experimental data. Since the model outcome and the experimental data are correlated, the scaling factor was calculated as the slope of the linear regression line between the two, with the formula given in Equation 11 (Eisenhauer, 2003).

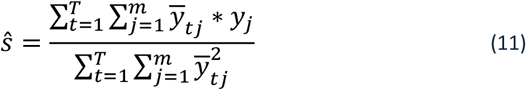

with *j* the number of strains, *y* is the set of experimental data point and 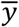 the model outcome.

The model of differential equations was solved with odeint from the scipy package in Python. The minimization was done with the lmfit package. As such, the parameters were all introduced with an initial estimate and search domain, being the minimal and maximal value. Next the cost function was minimized with the Nelder-Mead method by optimizing the parameter values. This cost function collected all the weighted residuals between the experimental and estimated data across all strains and variables (*i*.*e*., the mKate2/OD600 and sfGFP/OD600). The minimization method then searches the optimal parameters to reach the minimum of the Weighted Sum of Squared Residuals (*WSSR*(*θ*)) returned by the cost function, which is defined as follows:

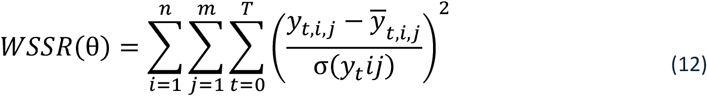

with *i* and *j* the number of variables and strains, respectively, y is the set of experimental data points, 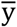 is the model outcome and θ the set of parameters. The residuals are weighted by the standard error across the three biological replicates (σ(*y*_*t*_*ij*)).

The obtained fits were further analyzed with a Markov Chain Monte Carlo (MCMC) analysis from the emcee package, incorporated in lmfit. The global sensitivity analysis was done with the sobol package from SALib.

## 5. Acknowledgments

This work was supported by the Research Foundation – Flanders (FWO) [grant numbers 1S29521N, 1246323N] and Special Research Fund (BOF) [grant number BOF20/IBF/131]. We would also like to thank the UGent Core Facility ‘HTS for Synthetic Biology for training, support and access to the instrument park. This Core Facility is supported by Special Research Fund (BOF) [grant numbers BOF/COR/2022/002, BAS018-18, BAS020-131 and BOF/BAS/2022/114] as well as FWO [grant numbers I011118N and I000925N].

## 6. Author contributions

J. De Baets designed and performed the experiments. J. De Baets analyzed the data and wrote the manuscript. B. De Paepe and M. De Mey were involved in the conceptualization and design of this work and in correcting and revising the manuscript.

## 8. Data availability

The datasets and computer code produced in this study are available in the following databases:

- Growth and fluorescence data: Github (https://github.com/MEMO-group/QuorumSensing_model_JDB.git)
- Modeling computer scripts: GitHub (https://github.com/MEMO-group/QuorumSensing_model_JDB.git)

